# PEXEL is a proteolytic maturation site for both exported and non-exported *Plasmodium* proteins

**DOI:** 10.1101/2023.07.12.548774

**Authors:** Manuel A Fierro, Ajla Muheljic, Jihui Sha, James A Wohlschlegel, Josh R Beck

## Abstract

Obligate intracellular malaria parasites dramatically remodel their erythrocyte host through effector protein export to create a niche for survival. Most exported proteins contain a pentameric *<u>P</u>lasmodium* <u>ex</u>port <u>el</u>ement (PEXEL)/Host Targeting Motif that is cleaved in the parasite ER by the aspartic protease Plasmepsin V (PMV). This processing event exposes a mature N-terminus required for translocation into the host cell and is not known to occur in non-exported proteins. Here we report that the non-exported parasitophorous vacuole protein UIS2 contains a *bona fide* PEXEL motif that is processed in the *P. falciparum* blood-stage. While the N-termini of exported proteins containing the PEXEL and immediately downstream ∼10 residues is sufficient to mediate translocation into the RBC, the equivalent UIS2 N-terminus does not promote export of a reporter. Curiously, the UIS2 PEXEL contains an unusual aspartic acid at the fourth position which constitutes the extreme N-terminal residue following PEXEL cleavage (P1’, RILτDE). Using a series of chimeric reporter fusions, we show that Asp at P1’ is permissive for PMV processing but abrogates export. Moreover, mutation of this single UIS2 residue to alanine enables export, reinforcing that the mature N-terminus mediates export, not PEXEL processing *per se*. Prompted by this observation, we further show that PEXEL sequences in the N-termini of other non-exported rhoptry proteins are also processed, suggesting that PMV may be a more general secretory maturase than previously appreciated, similar to orthologs in related apicomplexans. Our findings provide new insight into the unique N-terminal constraints that mark proteins for export.

**Importance:** Host erythrocyte remodeling by malaria parasite exported effector proteins is critical to parasite survival and disease pathogenesis. In the deadliest malaria parasite *Plasmodium falciparum*, most exported proteins undergo proteolytic maturation via recognition of the pentameric *<u>P</u>lasmodium* <u>ex</u>port <u>el</u>ement (PEXEL)/Host Targeting motif by the aspartic protease Plasmepsin V (PMV) which exposes a mature N-terminus that is conducive for export into the erythrocyte host cell. While PEXEL processing is considered a unique mark of exported proteins, we demonstrate PEXEL motifs are present and processed in non-exported proteins. Importantly, we show that specific residues at the variable fourth position of the PEXEL motif inhibit export despite being permissive for processing by PMV, reinforcing that features of the mature N-terminus, and not PEXEL cleavage, identify cargo for export cargo. This opens the door to further inquiry into the nature and evolution of the PEXEL motif.

## Introduction

*Plasmodium spp* are highly dependent on protein export for survival within the red blood cell (RBC) host. These proteins remodel the RBC to facilitate nutrient acquisition by the parasite and evasion of vertebrate host defenses (1). Proteins destined for export enter the secretory pathway via a signal sequence or transmembrane domain and are deposited into a vacuolar niche formed during invasion called the parasitophorous vacuole (PV) (2). Once in the PV, exported proteins are unfolded and transported across the PV membrane (PVM) via the PTEX complex to reach their final destination in the RBC space (3-7).

In *P. falciparum*, most exported proteins contain a *<u>P</u>lasmodium* <u>ex</u>port <u>el</u>ement (PEXEL,also known as the host-targeting signal), a pentameric motif (RxLxE/Q/D) generally located downstream of a recessed signal sequence (8, 9). ER entry occurs through a distinct SEC translocon complex that includes the SEC61 channel and SPC25, a non-catalytic component of the signal peptidase complex (10). During entry, the PEXEL sequence is recognized and cleaved between the third and fourth residues by Plasmepsin V (PMV), an aspartic protease that appears to operate in place of the classical signal peptidase SPC21 (11, 12). In keeping with ER translocation through a distinct SEC61 complex, the signal sequence of PEXEL proteins is generally not efficiently cleaved by signal peptidase and appears to act as a stable signal anchor that is removed through the action of PMV (13, 14). Following PMV cleavage, the newly exposed mature N-terminus is acetylated (15), although this widespread modification is not specific to exported proteins (16, 17) and is not sufficient for export (18).

PEXEL processing is crucial as export is blocked in non-cleaved mutants, which accumulate in the ER or PV (12, 18-22). Importantly, however, PMV processing can be bypassed by a reporter engineered to be processed by an exogenous protease to generate the equivalent of a mature PEXEL N-terminus (21). This indicates that PMV is not directly involved in cargo transfer in the export pathway but rather exposes an export-mediating mature N-terminus that is recognized by PTEX. While the mature protein retains only the last two PEXEL residues (xE/Q/D), the ∼10 residues immediately downstream of the PEXEL are also important for export but do not contain a discernable motif, suggesting that the N-terminal secondary structure adopted following processing may be important (19-21, 23). Furthermore, while PEXEL processing is necessary for export of proteins that contain this motif, it is not strictly required for translocation into the host cell as several <u>P</u>EXEL-<u>N</u>egative <u>E</u>xported <u>P</u>roteins (PNEPs) are known which also require PTEX translocation across the PVM but are not substrates for PMV (5, 6, 24). Indeed, despite the lack of PMV processing, the mature PNEP N-terminus is functionally equivalent to the mature PEXEL N-terminus in mediating export indicating that PEXEL processing is not strictly necessary for this process (19).

Although processing and export have not been validated for the full repertoire of PEXEL-containing proteins (20, 25-27), PEXEL cleavage is highly predictive for export and thought to be constrained to proteins that are trafficked into the erythrocyte. Interestingly, PEXEL processing has also been observed during the liver stage in proteins that are not exported beyond the PV into the hepatocyte (28, 29). As several of these proteins are also expressed in the blood stage and in some cases are translocated into the erythrocyte, this may correspond to a mechanistic difference in export between blood and liver stages; however, for the purple acid phosphatase-domain containing protein UIS2, localization also appears to be constrained to the PV during the blood stages (30, 31), suggesting a class of non-exported proteins exists that contain bona fide PEXEL motifs.

Here we show that although the UIS2 PEXEL is processed and N-acetylated in blood stage *P. falciparum,* the mature protein is not translocated beyond the PV, nor is the UIS2 leader sequence able to promote export of a fluorescent reporter despite PEXEL cleavage. Remarkably, a single amino acid change in the fourth position of the UIS2 PEXEL from aspartic acid to alanine enabled export of this reporter, showing that this variable residue which constitutes the first N-terminal amino acid after processing (P1’) is important for mediating export. Prompted by our findings with UIS2, we further identified PEXEL cleavage in several non-exported rhoptry proteins, some of which also contain a P1’ residue that restricts export. Collectively, our results indicate that PEXEL processing is a more general proteolytic maturation event than previously appreciated and reinforce that the mature N-terminus and not PEXEL processing *per se* identifies cargo for export. This work provides new insight into the unique N-terminal constraints that mark proteins for export and opens the door for further analysis of the nature and function of the PEXEL motif outside the context of export effectors.

## Results

### The PV protein UIS2 harbors a *bona fide* PEXEL motif that is not permissive for export

To query whether PEXEL processing extends to proteins that are not translocated into the RBC during intraerythrocytic development, we examined UIS2, a phosphatase domain-containing PV protein that encodes a classical signal peptide (residues 1-21) followed by a PEXEL motif (RILDE, residues 43-47) (Figure 1A) (30-32). Intriguingly, PEXEL processing of the *P. berghei* ortholog of UIS2 (RVLτQE) was detected by mass spectrometry analysis of liver-stage parasites (28) but has not been evaluated in the blood stage. To determine if the *P. falciparum* UIS2 PEXEL is also processed during the blood stages, we searched a previous mass spectrometry dataset that was generated from proximity labeling with a BioID2 fusion to the PV protein EXP2 in which UIS2 was well represented (Figure S1) (33). Indeed, peptide spectra corresponding to the processed, N-acetylated UIS2 PEXEL (cleavage after L45, Acetyl-DEYENINNSENEEDEYEDYLDDK) were identified, indicating similar PEXEL processing of *P. falciparum* UIS2 in the blood stage (Figure 1B). To ensure UIS2 was not exported beyond the PVM, we generated a C-terminal mNeonGreen (mNG) fusion to the endogenous copy of UIS2, which trafficked to the parasite periphery but was not observed in the erythrocyte (Figure 1C,D), consistent with strict localization to the PV as previously reported (30, 31).

**Figure 1.**
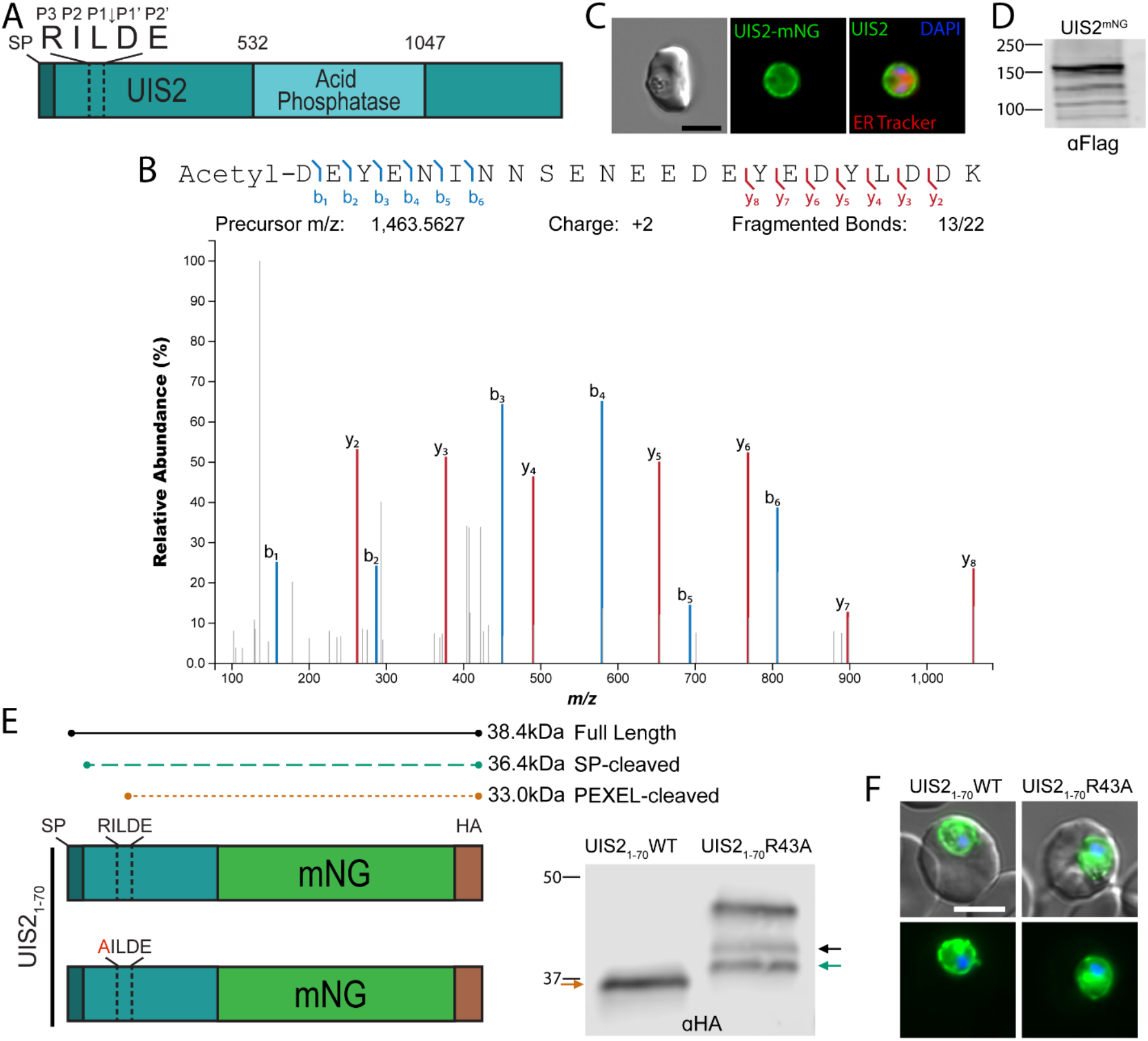
The PEXEL of UIS2 is cleaved but does not result in export. A) Schematic of full-length UIS2. Dark teal bar indicates signal peptide; dotted box indicates location of the PEXEL motif. B) Mass spectra of the N-terminal most UIS2 peptide detected by proximity labeling with EXP2-BioID2. The spectra correspond to PEXEL processing after L45 and acetylation at the new N-terminus. C) Representative live microscopy of endogenously tagged UIS2 with mNeonGreen (mNG) and 3XFlag tag and stained with ER-tracker. D) Western blot of UIS2-mNG-3xFlag parasites using anti-Flag antibodies. The expected size after PEXEL cleavage is 196 kDa. E) Schematic of reporter construct containing residues 1-70 of UIS2 with either the WT PEXEL motif or an R43A mutation fused to mNG and expressed under control of the *cam* promoter. Western blot of reporter constructs showing size differences indicating presence or absence of PEXEL cleavage. The expected molecular weights are labeled above the reporter schematic. The corresponding, colored arrows are shown on the gel image to denote the full length, signal peptidase cleaved or PEXEL cleaved versions of the reporters. F) Representative live microscopy of parasites expressing each reporter. Scale bar: 5µm.

The N-termini of PEXEL-containing exported proteins, including the PEXEL and immediately downstream spacer region (∼10 residues), are able to mediate export when fused to a fluorescent reporter protein (8, 9, 18, 34). To determine if more C-terminal features of UIS2 prevent its export, we fused the first 70 amino acids of UIS2, including the PEXEL and downstream 23 residues, to mNG with a C-terminal 3xHA epitope tag (Figure 1E). This cassette was placed under the control of the *cam* promoter and the plasmid was stably inserted into the *attB* site on chromosome 6 in NF54^attB^ parasites. Western blot analysis showed a single band at the expected size for PEXEL cleavage (Figure 1E, 1^st^ lane) and live fluorescence indicated this fusion protein was also secreted to the PV but failed to be exported into the erythrocyte (Figure 1F). To ensure PEXEL processing was still occurring in the context of the fusion protein, we generated a version where the PEXEL P3 arginine was changed to an alanine (R43A), which prevents processing by PMV (20). Consistent with previous reports (12, 18, 20), this mutation produced an upshift in molecular weight to the expected size for the signal peptidase cleaved version of the protein as well as a minor band that likely represents the full length, unprocessed form (Figure 1E, 2^nd^ lane, green and black arrows, respectively). An additional band that migrated at a higher molecular weight than the predicted size of the full-length protein was also observed in the mutated reporter, possibly indicating post-translational modification of residues N-terminal to the PEXEL motif. Localization of the R43A mutant was indistinguishable from the PEXEL processed fusion, trafficking to the PV but not the host cell (Figure 1F). Collectively, these results show that UIS2 contains a *bona fide* PEXEL motif but processing does not yield a mature N-terminus that can promote export, in contrast with other PEXEL containing exported proteins.

### UIS2 PEXEL position P1’ is not conducive for export

The ∼10-20 residues exposed upon PEXEL processing are critical for export, suggesting that the mature N-terminus of UIS2 lacks the necessary information to be recognized as cargo by PTEX. To determine if UIS2 residues immediately downstream of the PEXEL prevent export, we generated a mNG fusion reporter that replaced UIS2 residues 48-70 with residues 65-82 of the exported protein EMP3 previously shown to be sufficient for export in combination with the EMP3 PEXEL (Figure 2A, UIS2_1-47_-EMP3_65-82_) (20). In parallel, we placed a stretch of 12 alanine residues after the UIS2 PEXEL as this is also capable of mediating export when positioned immediately downstream of an export competent PEXEL (Figure 2C, UIS2_1-47_-Alanine_12_) (20). In both cases, we surprisingly observed full retention of the reporter at the PV with no export into the RBC (Figure 2A,C). To ensure this was not the result of an uncleaved PEXEL, we generated R43A mutant versions of these constructs and observed the expected upshift in molecular weight consistent with abrogation of PEXEL processing, confirming that the lack of export was not attributable to impaired UIS2 PEXEL processing in these chimeric constructs (Figure 2B,D).

**Figure 2.**
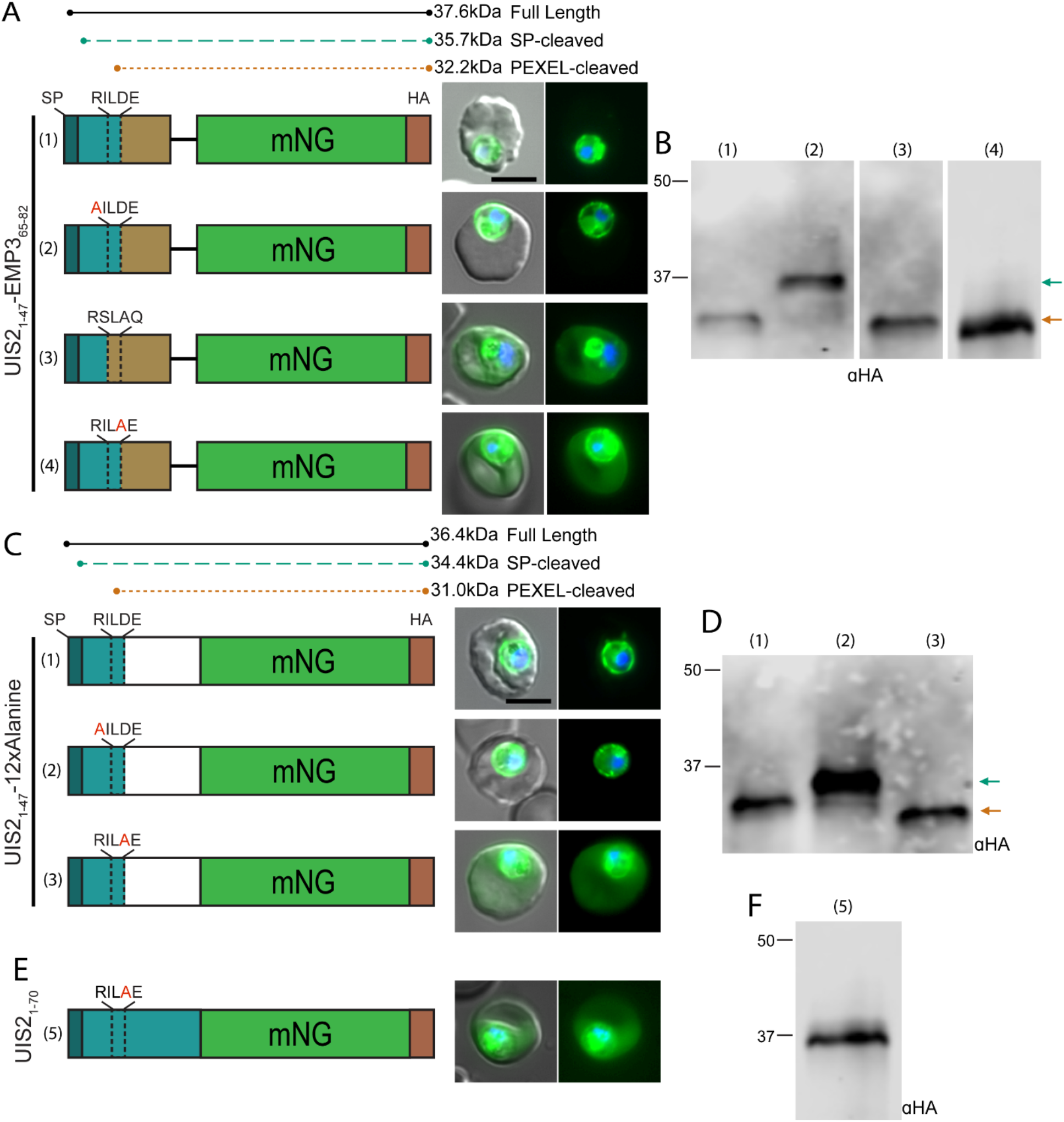
Aspartic acid at P1’ in the UIS2 PEXEL is not permissive for export. A) Schematic and representative live microscopy images of UIS2-EMP3 chimeric constructs containing residues 1-47 of UIS2 fused to residues 65-82 of EMP3 (1), the same construct containing R43A mutation of the UIS2 PEXEL (2), replacing the UIS2 PEXEL with that of EMP3 (3), or a R46A PEXEL mutation (4). B) Western blot of UIS2-EMP3 chimeric constructs. Arrows indicate bands corresponding to the expected molecular weights shown in the schematic in (A) associated with the presence or absence of PEXEL cleavage. C) Schematic and representative live microscopy images of UIS2-A12 chimeric constructs containing residues 1-47 of UIS2 fused to 12 alanine residues (1) and the same construct containing a R43A (2) or D46A mutation (3) of the UIS2 PEXEL. D) Western blot of UIS2-A12 chimeric constructs. Arrows indicate bands corresponding to the expected molecular weights shown in the schematic in (C) associated with the presence or absence of PEXEL cleavage. E) Schematic and representative live microscopy images of a construct containing residues 1-70 of UIS2 with an R43A mutation (5). F) Western blot of construct in (E). Scale bar: 5µm.

Lack of export in these chimeras indicates that either residues upstream of the UIS2 PEXEL or the PEXEL itself are inhibitory for export. To differentiate between these two possibilities, we replaced the UIS2 PEXEL with the PEXEL of EMP3 (RILDE>RLSAQ) in the UIS2-EMP3 chimera and found that this enabled export of mNG (Figure 2A,B). These results demonstrate that residues within the UIS2 PEXEL motif prohibit export, presumably amino acids at P1’ or P2’ that remain at the mature N-terminus. Previous reports have shown that mutations at the P1’ position of the PEXEL can inhibit export of chimeric reporters with minor impact on PEXEL processing (19, 21, 26). Amino acids with small, uncharged side chains such as serine, alanine and threonine are common at the P1’ position while the UIS2 PEXEL contains an unusual aspartic acid at P1’ which is not found in other predicted or identified exported proteins, suggesting this residue may not be compatible with an export-competent mature N-terminus (20, 26, 27). Indeed, replacing this aspartic acid with an alanine in both the UIS2_1-47_-EMP3_65-82_ and UIS2_1-47_-12xAlanine chimeras (Figure 2A,C) or the original UIS2 reporter (Figure 2E) enabled export. As expected, the D>A mutation at P1’ did not impact PEXEL processing of these construct (Figure 2B,D,F). These results show that aspartic acid at P1’ is permissive for PMV processing but not for export, reinforcing that the mature N-terminus, and not PEXEL processing *per se,* identifies cargo for export.

### Cleavable PEXEL motifs are present in other secreted proteins that do not traffic to the PV

Prompted by our observations with UIS2, we explored whether cleavable PEXELs were also present in other non-exported proteins. Prediction analyses have identified PEXEL motifs in rhoptry proteins which enter the secretory pathway but are not trafficked to the PV or exported into the host cell (20, 25-27). As such, we focused on the rhoptry-localized proteins RON11 and PMIX (Figure 3A,D) (35-37). RON11 is a large protein containing a classical signal peptide (residues 1-22) followed by an N-terminal PEXEL motif (RILFE, residues 51-55). This protein also contains 7 transmembrane domains and a single C-terminal EF-Hand motif (Figure 3A). Intriguingly, previous studies showed that the RON11 N-terminus containing the PEXEL was not conducive for export of a GFP reporter but export was enabled via an F54A mutation of the putative P1’ residue (25, 26). However, processing of these RON11 reporters was not evaluated. To determine if the RON11 PEXEL is cleaved, we fused wild type (WT) and R51A versions of residues 1-70 of RON11 to the mNG-3xHA reporter. The WT RON11 construct yielded a single band at the expected size for PEXEL processing while the R51A mutant produced upshifted bands with molecular weights corresponding to full-length and signal peptidase-cleaved species, indicating the RON11 PEXEL is cleaved (Figure 3C). Regardless of PEXEL processing, we observed a strict PV localization of both the control and R51A RON11 reporters with no evidence of export beyond the PV (Figure 3B). Similar to UIS2, mutating the RON11 P1’ residue to alanine (F54A, RILFE>RILAE) enabled export of mNG, demonstrating that phenylalanine at P1’ is compatible with PEXEL cleavage but also prohibits export (Figure 3B,C).

**Figure 3.**
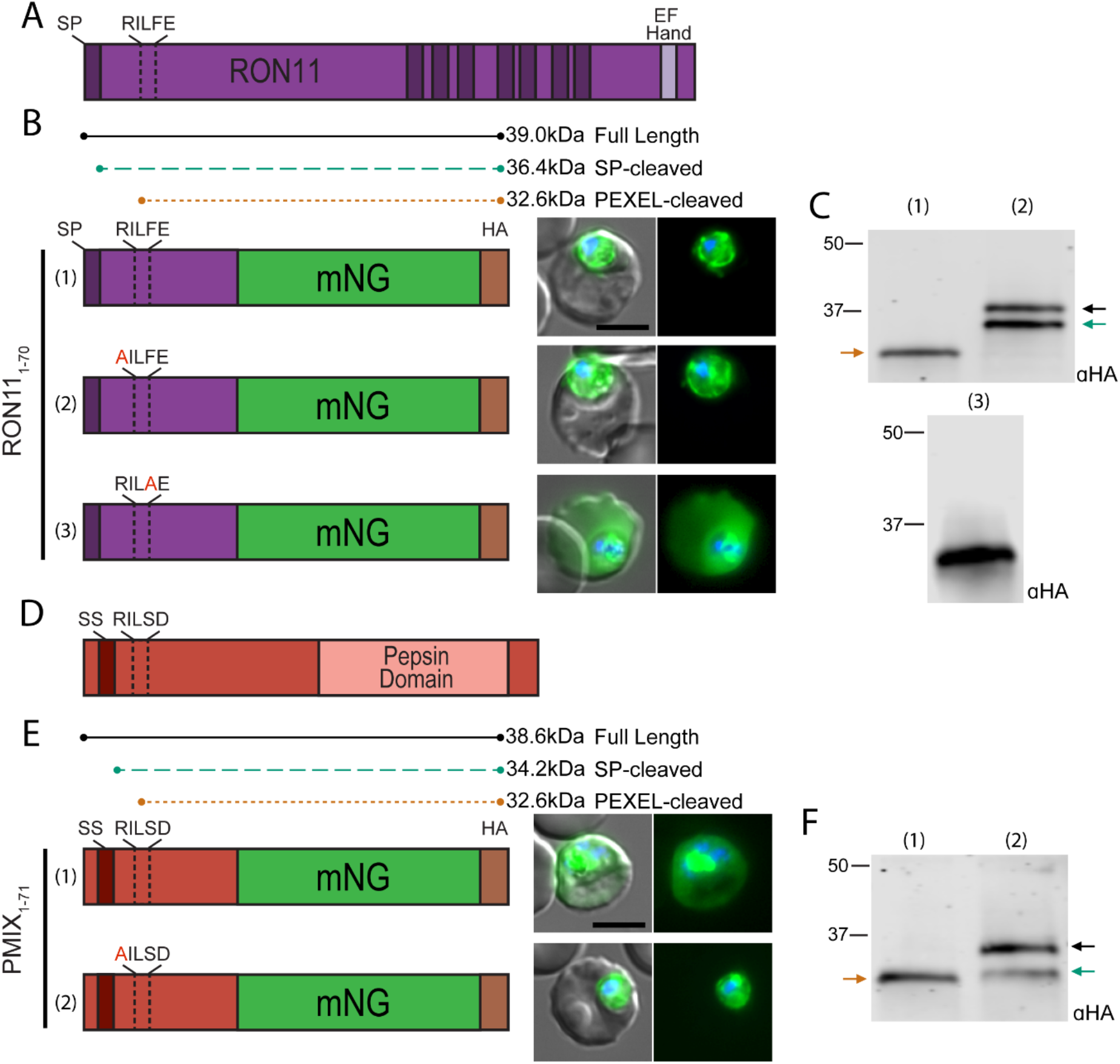
PEXEL motifs are present and cleaved in the N-termini of the rhoptry proteins RON11 and PMIX. A) Schematic of full-length RON11. Dark bars represent either the signal peptide or transmembrane domains. B) Schematic of reporter construct containing residues 1-70 of RON11 with either the WT PEXEL motif (1), an R51A (2) or F54A (3) PEXEL mutation, followed by representative live microscopy of parasites expressing the reporter. C) Western blot of RON11 reporter constructs. Arrows indicate bands corresponding to the expected molecular weights shown in the schematic in (B) associated with the presence or absence of PEXEL cleavage. D) Schematic of full-length PMIX. Dark red bar represents the recessed signal peptide. E) Schematic of reporter construct containing residues 1-71 of PMIX with either the WT PEXEL motif (1) or an R46A mutation (2) followed by representative live microscopy of parasites expressing the reporter. F) Western blot of PMIX reporter constructs. Arrows indicate bands corresponding to the expected molecular weights shown in the schematic in (E) associated with the presence or absence of PEXEL cleavage. Scale bar: 5µm.

We next examined the rhoptry protease Plasmepsin IX (PMIX) which contains an N-terminal hydrophobic sequence that may function as a classical signal peptide or recessed signal sequence (residues 1-39) followed by an N-terminal PEXEL (RILSD, residues 46-50) in a pro-region upstream of the aspartic peptidase domain (Figure 3D). Fusion of residues 1-71 of PMIX to an mNG reporter produced a single band at the expected size for the PEXEL cleaved species while an R46A mutation resulted in upshifted bands with molecular weights corresponding to full-length and signal peptidase-cleaved species, showing the PMIX PEXEL is processed in the blood stage (Figure 3F). Unlike the unusual aspartic acid or phenylalanine P1’ residues found in UIS2 or RON11, the PMIX PEXEL encodes a P1’ serine which is common among exported PEXEL proteins. Remarkably, the PMIX reporter fusion was exported, consistent with the importance of P1’ for determining the export competence of the mature PEXEL N-terminus. (Figure 3E). As expected, the R46A mutation eliminated export (Figure 3E). Taken together, these results show that PEXEL processing occurs not only in non-exported resident PV proteins but also in proteins that traffic to secretory organelles.

### Recessed signal sequence positioning in exported proteins is important for PEXEL processing

PEXEL-containing exported proteins are typically structured with an N-terminal hydrophobic sequence (sometimes referred to as “recessed signal sequence”) within a short first exon followed by the PEXEL motif encoded near the beginning of the second exon (8, 25). The recessed positioning of the signal sequence in exported proteins is thought to serve as a stable signal anchor and may be important for ER entry through a distinct SEC61 translocon complex that contains PMV in place of signal peptidase (10). In contrast, RON11 and UIS2 possess a canonical signal peptide positioned at their extreme N-terminus which is expected to mediate ER-entry and signal peptide processing through the classical SEC61 complex associated with the signal peptidase complex (Figure 1A and 3A) (10). The classical signal peptide configuration in UIS2 and RON11 suggests that the “recessed” signal sequence positioning in exported proteins may not be important for PEXEL processing and export. Indeed, replacing the EMP3 sequence upstream of the PEXEL (which contains a recessed signal sequence) with the equivalent region from UIS2 (which contains a signal peptide at the extreme N-terminus) supports export of mNG (Figure 2A), similar to previous observations indicating a recessed signal sequence is not necessary for PEXEL processing and export (20).

Since classical signal peptide and recessed signal sequence N-termini appear to be fully interchangeable for PEXEL processing, we tested whether a recessed signal sequence can function as a classical signal peptide when positioned at the extreme N-terminus. For this, we fused EMP3 residues 1-82 to mNG and compared this WT export reporter to a version where we removed residues 2-11 to place the hydrophobic stretch immediately after the start codon, mimicking the positioning of a classical signal peptide (Figure 4A). SignalP 3 (38) predicts a signal peptide for this arrangement with cleavage 24 residues upstream of the PEXEL, similar to the predicated signal peptidase cleavage sites for UIS2 and RON11, which are 21 and 28 residues upstream of the PEXEL, respectively. Interestingly, the truncated reporter was still exported but at noticeably lower levels compared to the WT control. Moreover, while the mNG signal within parasites expressing the WT reporter localized to the digestive vacuole (typical for proteins secreted to the PV or exported into the host cell which are partially endocytosed along with host cell cytosol), the mNG signal within parasites expressing the truncated reporter was distinctively perinuclear, suggesting that the repositioned signal sequence did not support ER exit (Figure 4A). Consistent with this, Western blot showed inefficient processing of the truncated reporter suggesting that the hydrophobic region of EMP3 is not recognized as a signal peptide and that positioning it at the N-terminus interferes with PEXEL processing (Figure 4B). Decreased export and increased ER retention was also observed with a similar truncation to reposition the signal sequence in the N-terminus of the PEXEL protein REX3 (Figure S2). Taken together, this indicates that the typical recessed position of the signal sequence in exported PEXEL proteins is important for PEXEL processing but is fully exchangeable with a classical signal peptide configuration at the extreme N-terminus for efficient PEXEL processing, suggesting that different modes of ER entry can support PEXEL maturation.

**Figure 4.**
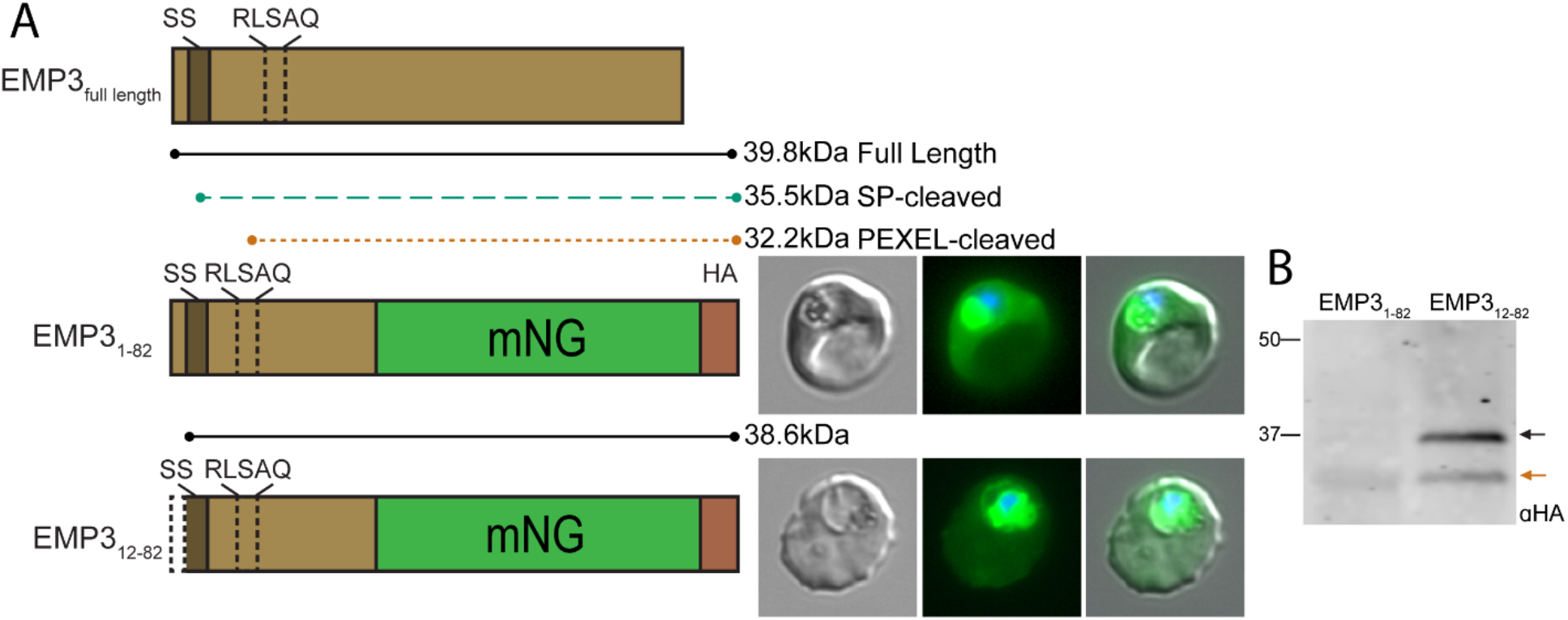
The position of the recessed signal sequence is important for PEXEL cleavage in EMP3. A) Schematic of full-length EMP3 accompanied by a schematic and representative live microscopy images of a reporter constructs containing the wild type recessed signal sequence or N-terminal truncation of residues 2-11 to place the signal sequence immediately after the start methionine. The signal within the parasite in the WT reporter line corresponds to the digestive vacuole, typical for proteins secreted to the PV or exported into the host cell, while signal within the parasite in the truncated reporter line is perinuclear. B) Western blot of wild type and truncated EMP3 constructs. Arrows indicate bands corresponding to the expected molecular weights shown in the schematic in (A) for full-length and PEXEL cleaved forms. Scale bar: 5µm.

## Discussion

The discovery of the PEXEL motif revolutionized the identification of host targeted effector proteins in *Plasmodium* due to its high predictive power for export (8, 9, 20, 25-27). Consequently, PEXEL processing is generally thought to be restricted to exported proteins. In this study, we reveal PEXEL processing also occurs in the non-exported PV and rhoptry proteins UIS2, RON11, and PMIX. While deviations at the strongly constrained P3, P1 and P2’ positions have been shown to support processing and export in certain contexts (27), the flexibility of residues occupying these positions is limited (14, 18, 20-22, 39). In contrast, a wide range of charge-neutral residues can occupy P2 and P1’. While successful PEXEL cleavage generally leads to protein export, the P1’ residue, which becomes the N-terminal most position in the mature N-terminus, can interfere with export after processing (19, 21, 26). Remarkably, we found that the inability of the mature N-termini of UIS2 and RON11 to mediate export is attributable to this single amino acid (aspartic acid or phenylalanine) at position P1’ in the PEXEL of these proteins. Notably, the UIS2 PEXEL motif differs between *P. berghei* and *P. falciparum* (RVLQE vs. RILDE), indicating that glutamine at P1’ likely also inhibits export. Thus PEXEL motifs with charged or bulky residues at P1’ are competent for processing but not export. Consistent with this, a recent report indicates lysine is also not tolerated for export at P1’ (40). For PV resident proteins that do not function in the host cell, these P1’ residues would provide a control mechanism to allow for proteolytic maturation by PMV without risking unproductive export, presumably preventing recognition by the PTEX unfoldase HSP101 (41).

We also found that PEXEL motifs are processed in the rhoptry-localized proteins RON11 and PMIX, indicating that PEXEL processing is a more general maturation event than previously appreciated. While the P1’ of RON11 prevents export, similar to UIS2, PMIX encodes a serine at position P1’ which is found in the PEXEL of many exported proteins (8, 9, 25) and processing of the PMIX PEXEL did result in reporter export. Exported proteins follow the default secretory pathway to the PV while proteins that traffic to other destinations contain targeting information that overrides default PV secretion (42, 43). This suggests that the dominant rhoptry trafficking signal in PMIX prevents it reaching the PV and being exported. While unfolding and PVM translocation of exported proteins occurs in the vacuole, it should be noted that HSP101 has recently been proposed to recognize cargo upstream in the ER (22). However, PMIX undergoes an additional autocatalytic maturation step (p73>p55) that removes ∼100 residues after the PEXEL (35, 36, 44), eliminating the mature PEXEL N-terminus that would otherwise drive export of the protease. As no N-terminal tagging has been applied to PMIX, it is not clear whether this cleaved fragment is ultimately exported beyond the parasite. Notably, while the RON11 PEXEL is widely conserved in orthologs from other *Plasmodium* spp, the PMIX PEXEL is not (Figure S3). The functional significance of PEXEL processing in PfPMIX and closely related *Laverania* species is unclear but lack of PEXEL conservation implies that initial cleavage by PMV is not generally important for activating PMIX to enable its ultimate autocatalytic maturation.

Entry into the secretory pathway via a signal peptide or transmembrane domain is mediated by the SEC61 complex which, in *Plasmodium*, appears to form distinct associations with either signal peptidase or PMV (10). While exported PEXEL proteins typically contain a recessed signal sequence thought to act as a membrane anchor that is removed by PMV, UIS2 and RON11 encode a classical signal peptide suggesting that multiple ER entry points are tolerated enroute to PMV. If that were the case, then repositioning the recessed signal sequence domain of exported proteins in a conformation that mimics a classical signal peptide should still route these proteins to PMV through the signal peptidase-SEC61 complex. Indeed, we observed a portion of the reporter was still exported, indicative of processing by PMV. However, repositioning of the recessed signal sequence to the extreme N-terminus resulted in increased ER retention of EMP3 and REX3, indicating that the repositioned signal sequence mediates insertion into the ER membrane but is not efficiently recognized and cleaved by either signal peptidase or PMV. Indeed, several PEXEL mutants that block PMV processing accumulate as the unprocessed, full-length form of the protein, showing that the recessed signal sequence is not efficiently recognized as a signal peptide (10, 12-14, 18, 20, 27). Surprisingly, repositioning the signal sequence in EMP3 caused a major reduction in PEXEL processing. As this construct preserves the spacing between the signal sequence and PEXEL that is important for cleavage by PMV (14), inefficient processing could indicate an altered topology where the truncated protein is inserted into the ER membrane as a type III transmembrane protein, placing the C-terminus in the cytoplasm where it is inaccessible to PMV (45). Curiously, P3 R>A mutations that blocked PEXEL processing in RON11 and UIS2 accumulated as two distinct species, suggesting that the signal peptide in these proteins is not efficiently recognized by signal peptidase and may act as a stable signal anchor similar to PEXEL proteins encoding recessed signal sequences. If this is the case, then PEXEL-containing proteins may enter the ER through the PMV-SEC61 complex regardless of signal sequence positioning.

The discovery of PEXEL processing in non-exported proteins may indicate that PMV was historically a more general secretory maturase that subsequently became overly dedicated to exported proteins through expansion of exported proteins families, such as RIFINs and PHISTs in *P. falciparum* (25). As the parasite became heavily dependent on protein export, the PMV-SEC61 complex may have been adapted to alleviate the burden on the canonical SEC61 complex. Of note, a similar *Toxoplasma* Export Element (TEXEL, RRLxx) is cleaved by the PMV ortholog ASP5. Although TEXEL processing is also important for vacuolar export of *Toxoplasma* effector proteins (46-48), many ASP5 substrates are PV-resident proteins that are not translocated into the host cell (49) and processing has been shown to be dispensable for localization and function of some non-exported proteins (50). In apparent contrast to *Plasmodium*, TEXEL processing is not restricted to the N-terminus and some proteins are processed at multiple TEXELs, such as the UIS2 ortholog GRA44 (50-52). C-terminally located PEXEL sequences can also be found in *Plasmodium* proteins but are not known to be processed. For instance, ClpB1, a nuclear encoded HSP100 that is targeted to the apicoplast, contains an identical PEXEL to PfUIS2 (RILDE) located at positions 1044-1048 of 1070; however, this does not appear to be appreciably processed as C-terminal epitope tags are retained on the full length protein, even when it is artificially retrieved to the ER (53).

Curiously, ASP5 is located in the Golgi (54) where post-translational modifications in eukaryotes typically occur, suggesting that export cargo recognition functions differently in *T. gondii*. Indeed, *T. gondii* possess an analogous but mechanistically different vacuolar export pathway that depends upon the MYR complex thought to form a distinct translocon (52, 55, 56). That PEXEL processing occurs coincidently with ER entry in *Plasmodium* suggests that the mature, export mediating N-terminus may be recognized early in the secretory pathway, presumably by HSP101, although it is unclear why this should occur upstream of the site of membrane translocation by PTEX (22). Future studies that determine why certain mature PEXEL N-termini enable selection for PVM translocation will be important to resolve the connection between these key events at opposite ends of the parasite secretory pathway.

## Methods

### Parasite culture

*P. falciparum* NF54^attB-DiCre^ (53) and derivatives were cultured in RPMI 1640 medium supplemented with 27 mM sodium bicarbonate, 11 mM glucose, 0.37 mM hypoxanthine, 10 µg/ml gentamicin and 0.5% Albumax I (Gibco). Parasites were maintained in deidentified, Institutional Review Board-exempt RBCs obtained from the American National Red Cross.

### Plasmids and genetic modification of *P. falciparum*

All cloning was carried out with NEBuilder HIFI DNA assembly (NEB). Primer sequences are given in Table S1. For Cpf1-mediated editing of the UIS2 locus, the pAIO-LbCpf1 plasmid (33) was modified by inserting a HindIII site adjacent to the AflII site in the pre-gRNA cassette using primer P1, resulting in the plasmid JRB489, which enables double digestion to reduce parent vector background during cloning. To target the 3’ end of *uis2*, a Cpf1 gRNA target was chosen just downstream of the stop codon (TATTACCTTGATATTCTATTAAGG) and the gRNA seed sequence was synthesized in primer P2 and inserted at HindIII/AflII in JRB489, resulting in plasmid JRB484. To generate UIS2-mNG parasites, a 5’ homology flank (up to but not including the stop codon) and 3’ homology flank (beginning immediately downstream of the Cpf1 gRNA PAM) were amplified from genomic DNA using P3/4 and P5/6, respectively, assembled in a second PCR reaction using P4/5 and inserted between XhoI/AvrII in JRB508 (53), resulting in plasmid JRB509. This plasmid was linearized with AflII and co-transfected with the corresponding JRB484 into NF54^attB-DiCre^. Selection was applied with 5 nM WR99120 24 hrs post-transfection and clonal lines were isolated by limiting dilution after parasites returned from selection.

To generate the UIS2_1-70_-mNG fusion cassette, the 5’ cds of *uis2* encoding the first 70 amino acids was amplified from gDNA using P7/8 and inserted at AvrII in JRB416 (53) removing the adjacent loxP site and inserting an NheI site between the *uis2* and mNG sequences, resulting in plasmid JRB510. A R43A PEXEL mutation in UIS2_1-70_-mNG was generated in this plasmid using a QuikChange Lightning Multi Site Directed Mutagenesis kit (Agilent) and the primer P9, resulting in plasmid JRB525 while a D45A PEXEL mutation was done using primers P7/32 and P33/7 to generate two PCR products that were then stitched together in a second PCR reaction using primers P7/8 resulting in plasmid MAF63. To generate the UIS2-EMP3 chimeric constructs, primer pairs were used to amplify the leader sequence of UIS2 including the endogenous PEXEL (P7/P10), a R43A PEXEL mutation (P7/P11), a D45A PEXEL mutation (P7/4), and replacing the UIS2 PEXEL with the EMP3 PEXEL (P7/P12), while appending residues 65-74 of EMP3. A second PCR reaction was done using this initial PCR as a template using primers P7/13 to append residues 75-82 of EMP3 which was subsequently inserted into JRB510 between AvrII/NheI generating plasmids MAF45, MAF52, MAF61, MAF53, respectively. A similar approach was used to generate UIS2-A12 chimeras where primer pairs were used to amplify the leader sequence of UIS2 including the endogenous PEXEL (P7/P14), R43A PEXEL mutation (P7/P15), and a R46A PEXEL mutation (P7/P16), while appending 12 Alanine residues after the PEXEL. These were inserted into JRB510 between AvrII/NheI generating plasmids MAF48, MAF59, MAF57, respectively. These plasmids were co-transfected with pINT (57) into NF54^attB-DiCre^ parasites and selection was applied with 2.5 µg/ml blasticidin-S 48 hrs post-transfection.

To generate the EMP3-mNG fusion construct, primers P17/P18/P19/P13 were used to synthetically generate residues 1-82 of EMP3. Using 10µM of each primer, an initial PCR reaction was performed using the following parameters: 94°C for 5min, [94°C for 2min, 60°C for 2min, 68°C for 3min] x 9 rounds, 68°C for 5min, 4°C ∞. One µL of this initial reaction was then used in a subsequent, standard PCR reaction using primers P17/P13 which was then inserted into JRB510 between AvrII/NheI generating plasmid MAF51. To generate EMP3-mNG without the recessed signal peptide, a similar approach was used except using primers P20/P18/P19/P13 in an initial PCR reaction and then using primers P20/P13 to generate the final product inserted into JRB510 between AvrII/NheI generating plasmid MAF54.To generate the REX3-mNG fusion construct, primers P21/P22 were used to amplify residues 1-70 of REX3 while primers P23/P22 were used to amplify REX3 without the recessed signal peptide. These plasmids were co-transfected with pINT into NF54^attB-DiCre^ parasites and selection was applied with 2.5 µg/ml blasticidin-S 48 hrs post-transfection.

To generate the RON11-mNG fusion cassette, primers P24/P25 were used to amplify residues 1-70 of RON11, while removing the N-terminal intron, and inserted into JRB510 between AvrII/NheI generating plasmid MAF47. To generate the R51A or F54A mutation of the RON11 PEXEL, plasmid MAF47 was linearized with the site BstBI that resides within the PEXEL and primer P26 or P35, containing the PEXEL mutation, was used to seal the vector using NEBuilder HIFI DNA assembly resulting in plasmids MAF50 and MAF60. To generate the RON11-EMP3 chimera containing the F54A PEXEL mutation, primers P24/36 were used to append residues 65-74 of EMP3. A second PCR reaction was done using this initial PCR as a template using primers P24/13 to append residues 75-82 of EMP3 generating plasmid MAF62. To generate the PMIX-mNG fusion cassette, primers P27/P28 were used to amplify residues 1-71 of PMIX and inserted into JRB510 between AvrII/NheI generating plasmid MAF46. To generate the R46A PEXEL mutation, primers P27/P29/P30/P31 were used to synthetically generate residues 1-71 of PMIX with the PEXEL mutation in an initial PCR reaction, as described above, followed by a second PCR reaction using primers P27/P28, which was then inserted into JRB510 between AvrII/NheI generating plasmid MAF49.

### Microscopy

For live microscopy, Hoescht (33342, Invitrogen; 1:10,000) was used to visualize the nucleus by incubating with parasites 2-5min prior to visualization. ER-Tracker Red (Invitrogen; 1:1000) was used to visualize the ER by incubating with parasites 30min at 37°C prior to visualization. Parasites were imaged on an Axio Observer 7 equipped with an Axiocam 702 mono camera and Zen 2.6 Pro software (Zeiss) using the same exposure times for all images across sample groups and experimental replicates. Image processing, analysis, and display were performed using Zeiss Blue software and Adobe Photoshop. Equivalent adjustments to brightness and contrast were made within sample groups for display purposes.

### Western blotting

Western blotting for *Plasmodium* parasites was performed as described previously (33, 58). Briefly, parasites were selectively permeabilized by treatment with ice-cold 0.03% saponin in PBS for 15 min followed by lysis with RIPA to remove the hemozoin. The antibodies used in this study were mouse anti-HA antibody (clone 16B12, BioLegend; 1:500) and rabbit polyclonal anti-HA SG77 (ThermoFisher; 1:500). The secondary antibodies used are IRDye 680CW goat anti-mouse IgG (LICOR Biosciences) (1:20,000). Western blot images were processed using the Odyssey Clx LICOR infrared imaging system software (LICOR Biosciences). Full-length blots are presented in Figure S4.

### LC-MS Acquisition and Analysis

For sample preparation of parasite lysates, see (33). Peptide samples were separated on a 75uM ID x 25cm C18 column packed with 1.9μm C18 particles (Dr. Maisch GmbH) using on a 140-minute gradient of increasing acetonitrile and eluted directly into a Thermo Orbitrap Fusion Lumos mass spectrometer where MS spectra were acquired by Data Dependent Acquisition (DDA). Data were searched using MaxQuant and a customized *Plasmodium falciparum 3D7* database containing the predicted N-terminally processed version of PF3D7_1464600 with carbamidomethylation set as a fixed modification and methionine oxidation and N-term acetylation set as variable modifications. The digestion mode was set to trypsin with a maximum of 2 missed cleavages. Precursor mass tolerances of 20 ppm and 4.5 ppm were applied to the first and second searches, respectively, while a 20-ppm mass tolerance was used for fragment ions. The datasets were filtered at a 1% false discovery rate (FDR) at both the peptide-spectrum match (PSM) and protein levels.

## Acknowledgements

This study was supported by NIH grant AI173810 to JRB. The funders had no role in study design, data collection and interpretation, or the decision to submit the work for publication.

## Author contributions

MAF and JRB conceived and designed the experiments. MAF, AM, JS, JW and JRB performed the experiments. MAF, AM, JS, JW and JRB analyzed the data. MAF and JRB wrote the manuscript. All authors discussed and edited the manuscript.

---

**Figure S1.**
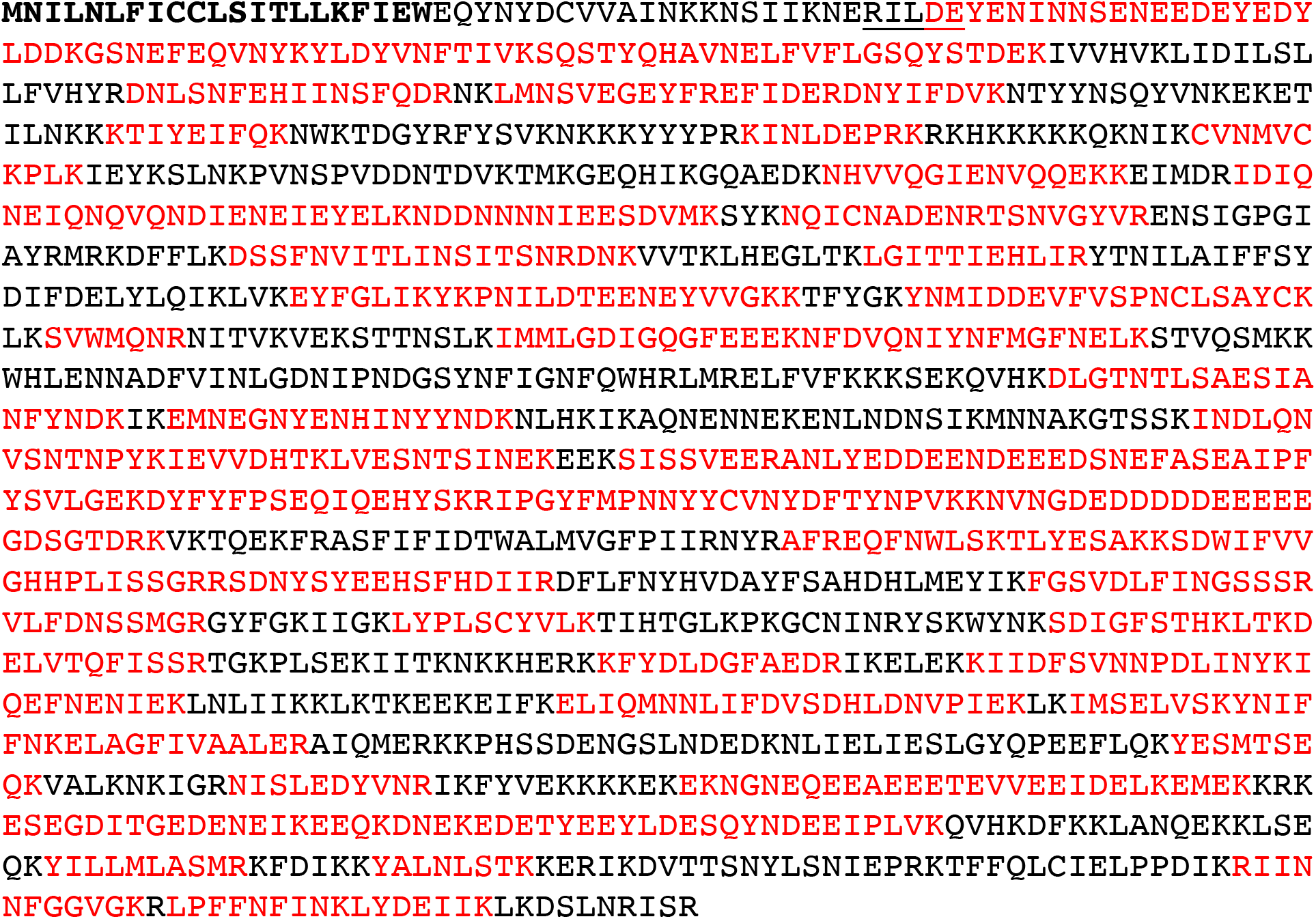
UIS2 peptides identified by LC/MS-MS from EXP2-BioID2 proximity labeling experiments (33). Peptides are highlighted in red. The PEXEL is underlined and the signal peptide predicted by SignalP 4.1 (59) is shown in bold. The N-terminal most peptide identified corresponds to the processed and N-acetylated PEXEL (see Figure 1B).

**Figure S2.**
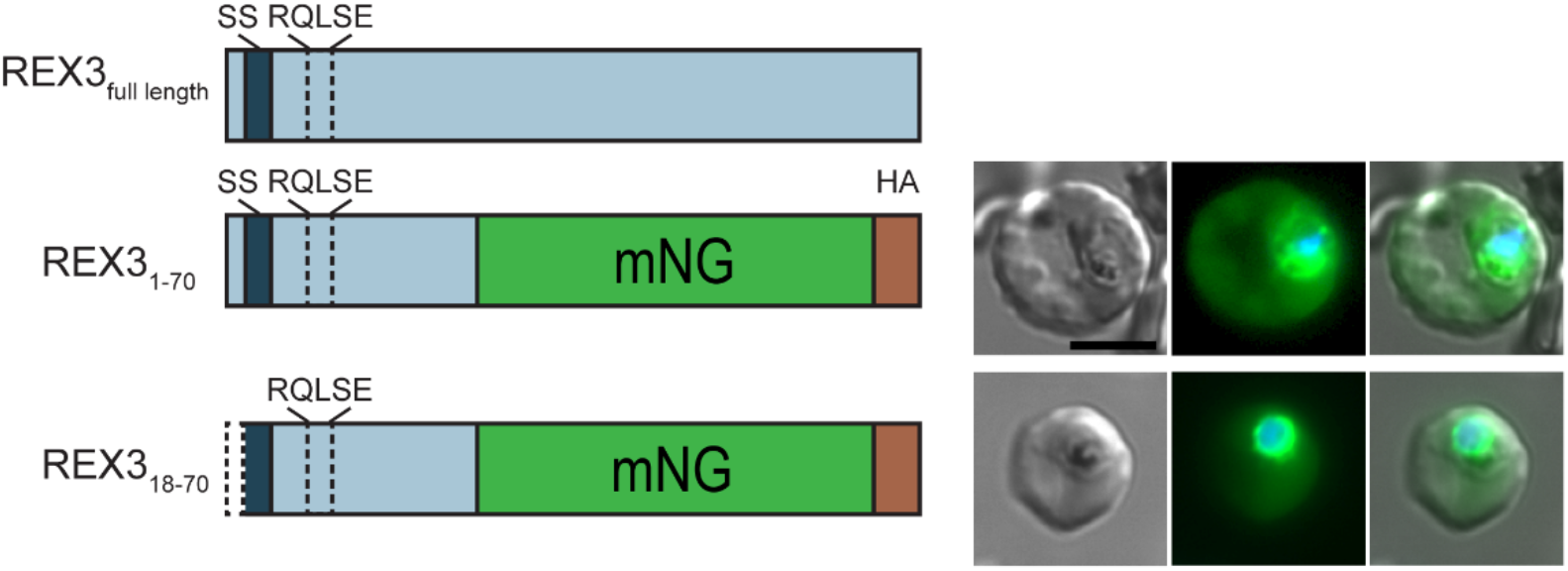
Schematic of full-length REX3 (top) and schematics and representative live microscopy images of reporter constructs containing the wild type recessed signal sequence or N-terminal truncation removing residues 2-17 to place the signal sequence immediately after the start methionine to mimic a classical signal peptide. SignalP 3 (38) predicts a signal peptide for this arrangement with cleavage 14 residues upstream of the PEXEL. Scale bar: 5µm.

**Supplementary Figure 3.**
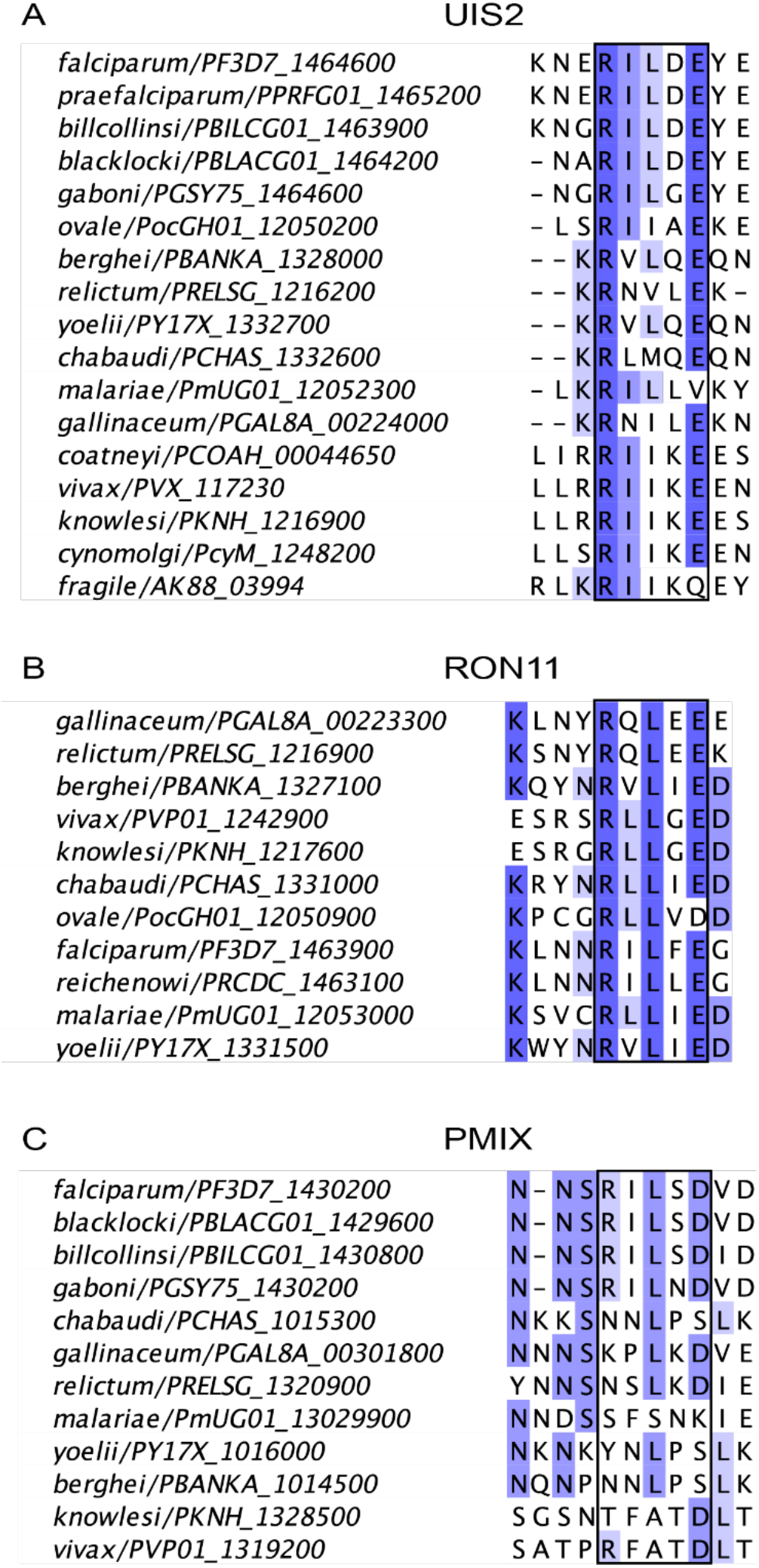
Alignment of (A) UIS2, (B) RON11 or (C) PMIX ortholog sequences from indicated *Plasmodium* spp (species/gene number). Alignments were generated using Clustal Omega (60) and a portion of the alignment corresponding to a 10 amino acid window containing the PEXEL motif from each *P. falciparum* ortholog was then displayed using Jalview (61). A) Canonical PEXEL sequences are present in UIS2 from *Laverania* species, *P. berghei* and *P. yoelii* while most other species encode a non-canonical PEXEL (RxIxE/D/Q) known to be processed in some contexts (27). P1’ residues across all canonical and non-canonical variants are expected to be non-permissive for export (D, Q and K) except for *P. gaboni* (RILGE), *P. ovale* (RIIAE) and *P. gallinaceum* (RNILE). B) Canonical PEXEL sequences are broadly conserved in RON11 although some encode P1’ residues expected to be permissive to export. C) Canonical PEXEL sequences are not present in PMIX orthologs outside of *Laverania* species. A non-canonical PEXEL (KxLxE/Q/D) is present in *P. gallinaceum* (KPLKD).

**Supplementary Figure 4.**
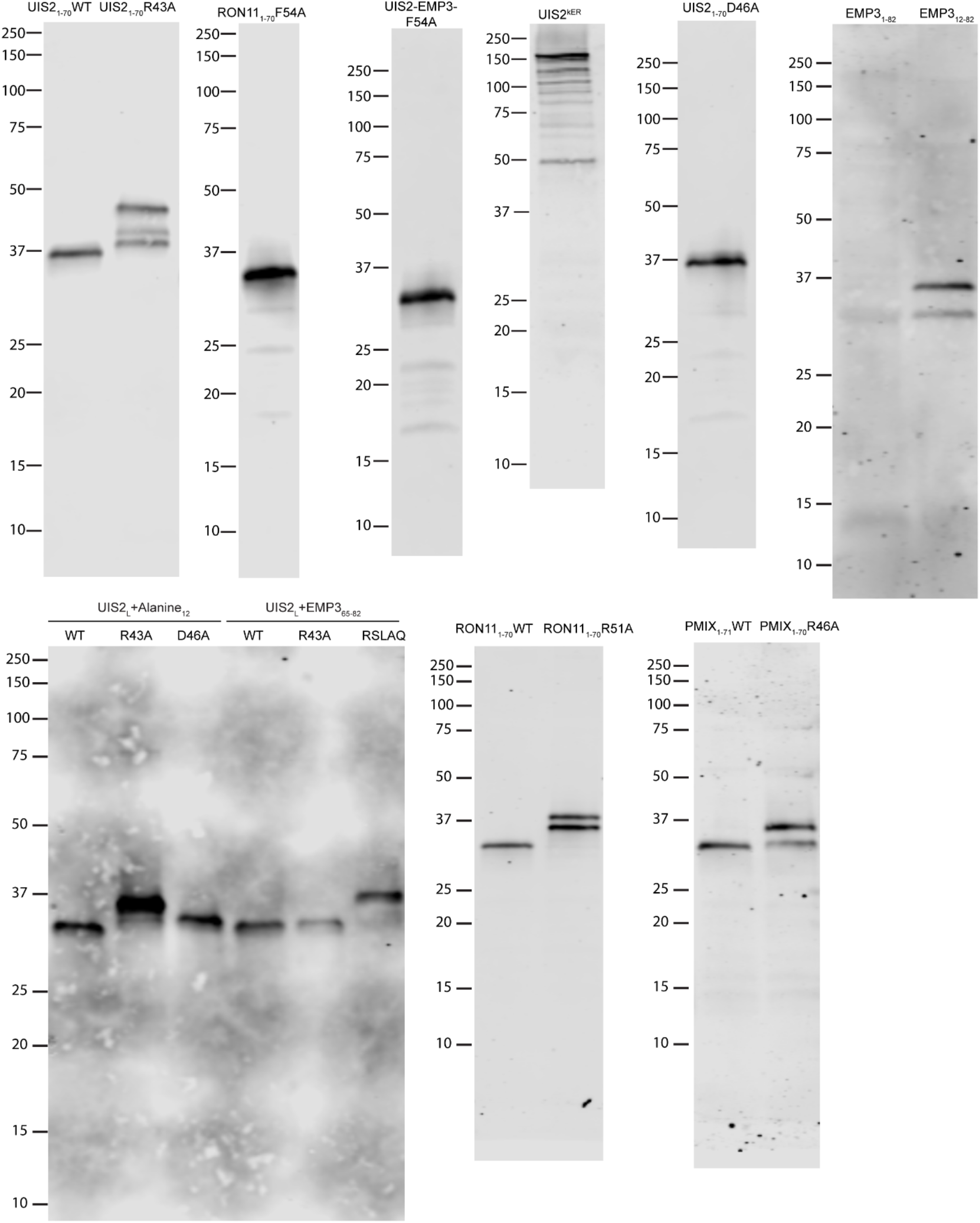
Uncropped western blots from this study.

**Table S1.**
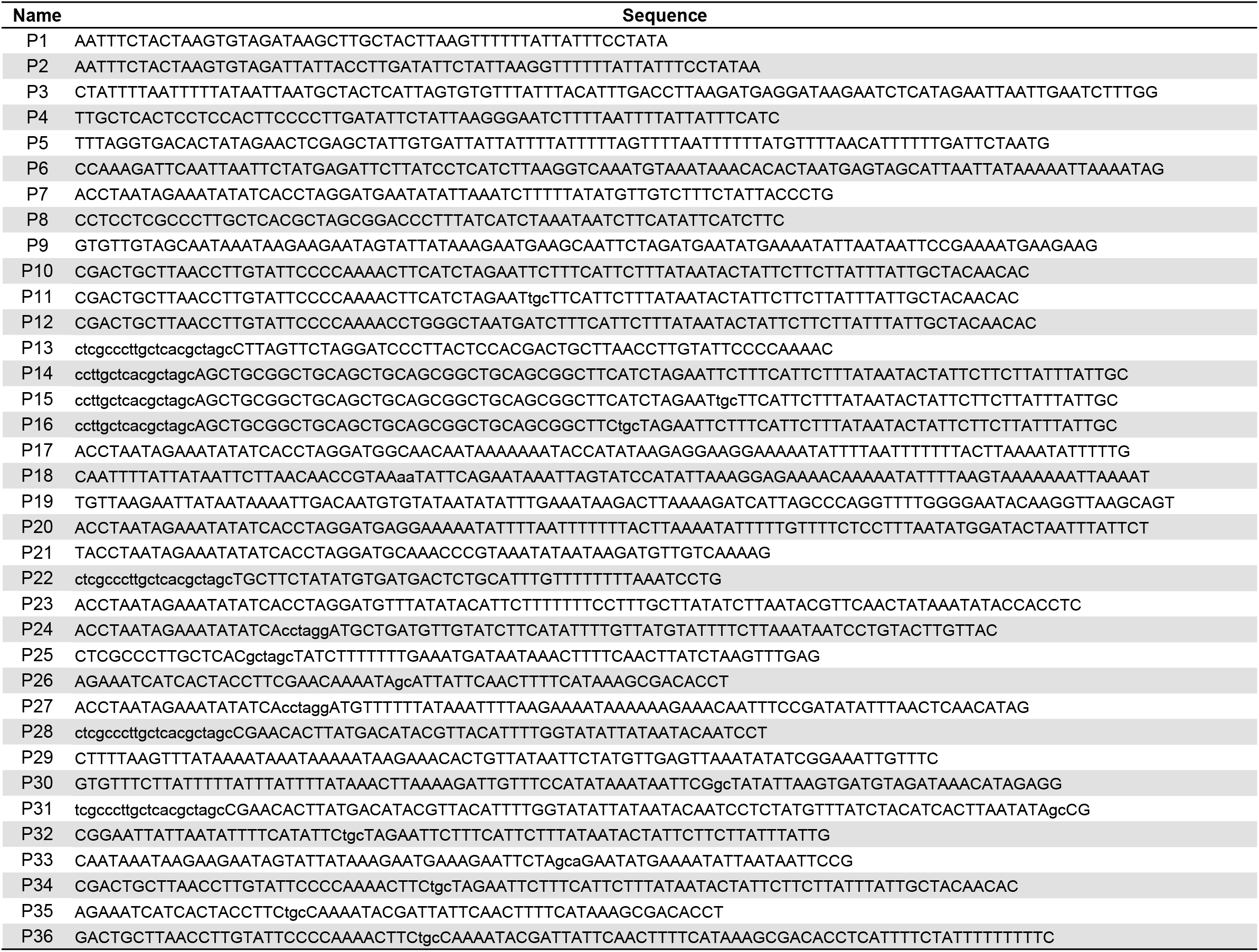
Sequences of primers used in this study.

## References

1. T. K. Jonsdottir, M. Gabriela, B. S. Crabb, T. F. de Koning-Ward, P. R. Gilson, Defining the Essential Exportome of the Malaria Parasite. Trends Parasitol 37, 664–675 (2021).

2. J. M. Matz, J. R. Beck, M. J. Blackman, The parasitophorous vacuole of the blood-stage malaria parasite. Nat Rev Microbiol 18, 379–391 (2020).

3. N. Gehde et al., Protein unfolding is an essential requirement for transport across the parasitophorous vacuolar membrane of Plasmodium falciparum. Mol Microbiol 71, 613–628 (2009).

4. T. F. de Koning-Ward et al., A newly discovered protein export machine in malaria parasites. Nature 459, 945–949 (2009).

5. J. R. Beck, V. Muralidharan, A. Oksman, D. E. Goldberg, PTEX component HSP101 mediates export of diverse malaria effectors into host erythrocytes. Nature 511, 592–595 (2014).

6. B. Elsworth et al., PTEX is an essential nexus for protein export in malaria parasites. Nature 511, 587–591 (2014).

7. P. Mesen-Ramirez et al., Stable Translocation Intermediates Jam Global Protein Export in Plasmodium falciparum Parasites and Link the PTEX Component EXP2 with Translocation Activity. PLoS Pathog 12, e1005618 (2016).

8. M. Marti, R. T. Good, M. Rug, E. Knuepfer, A. F. Cowman, Targeting malaria virulence and remodeling proteins to the host erythrocyte. Science 306, 1930–1933 (2004).

9. N. L. Hiller et al., A host-targeting signal in virulence proteins reveals a secretome in malarial infection. Science 306, 1934–1937 (2004).

10. D. S. Marapana et al., Plasmepsin V cleaves malaria effector proteins in a distinct endoplasmic reticulum translocation interactome for export to the erythrocyte. Nat Microbiol 3, 1010–1022 (2018).

11. I. Russo et al., Plasmepsin V licenses Plasmodium proteins for export into the host erythrocyte. Nature 463, 632–636 (2010).

12. J. A. Boddey et al., An aspartyl protease directs malaria effector proteins to the host cell. Nature 463, 627–631 (2010).

13. B. E. Sleebs et al., Inhibition of Plasmepsin V activity demonstrates its essential role in protein export, PfEMP1 display, and survival of malaria parasites. PLoS Biol 12, e1001897 (2014).

14. J. A. Boddey et al., Export of malaria proteins requires co-translational processing of the PEXEL motif independent of phosphatidylinositol-3-phosphate binding. Nat Commun 7, 10470 (2016).

15. H. H. Chang et al., N-terminal processing of proteins exported by malaria parasites. Mol Biochem Parasitol 160, 107–115 (2008).

16. M. A. Nyonda et al., N-acetylation of secreted proteins in Apicomplexa is widespread and is independent of the ER acetyl-CoA transporter AT1. J Cell Sci 135 (2022).

17. A. J. Polino et al., An essential endoplasmic reticulum-resident N-acetyltransferase ortholog in Plasmodium falciparum. J Cell Sci 136 (2023).

18. J. A. Boddey, R. L. Moritz, R. J. Simpson, A. F. Cowman, Role of the Plasmodium export element in trafficking parasite proteins to the infected erythrocyte. Traffic 10, 285–299 (2009).

19. C. Gruring et al., Uncovering common principles in protein export of malaria parasites. Cell Host Microbe 12, 717–729 (2012).

20. J. A. Boddey et al., Role of plasmepsin V in export of diverse protein families from the Plasmodium falciparum exportome. Traffic 14, 532–550 (2013).

21. S. J. Tarr, A. Cryar, K. Thalassinos, K. Haldar, A. R. Osborne, The C-terminal portion of the cleaved HT motif is necessary and sufficient to mediate export of proteins from the malaria parasite into its host cell. Mol Microbiol 87, 835–850 (2013).

22. M. Gabriela et al., A revised mechanism for how Plasmodium falciparum recruits and exports proteins into its erythrocytic host cell. PLoS Pathog 18, e1009977 (2022).

23. S. Haase et al., Sequence requirements for the export of the Plasmodium falciparum Maurer’s clefts protein REX2. Mol Microbiol 71, 1003–1017 (2009).

24. A. Heiber et al., Identification of New PNEPs Indicates a Substantial Non-PEXEL Exportome and Underpins Common Features in Plasmodium falciparum Protein Export. PLoS Pathog 9, e1003546 (2013).

25. T. J. Sargeant et al., Lineage-specific expansion of proteins exported to erythrocytes in malaria parasites. Genome Biol 7, R12 (2006).

26. C. van Ooij et al., The malaria secretome: from algorithms to essential function in blood stage infection. PLoS Pathog 4, e1000084 (2008).

27. J. Schulze et al., The Plasmodium falciparum exportome contains non-canonical PEXEL/HT proteins. Mol Microbiol 97, 301–314 (2015).

28. M. J. Shears et al., Proteomic Analysis of Plasmodium Merosomes: The Link between Liver and Blood Stages in Malaria. J Proteome Res 18, 3404–3418 (2019).

29. R. McConville et al. (2023) Plasmodium falciparum exoerythrocytic forms require the PTEX translocon for development in human hepatocytes. (Research Square Platform LLC).

30. M. Khosh-Naucke et al., Identification of novel parasitophorous vacuole proteins in P. falciparum parasites using BioID. Int J Med Microbiol 308, 13–24 (2018).

31. C. B. Schnider, D. Bausch-Fluck, F. Bruhlmann, V. T. Heussler, P. C. Burda, BioID Reveals Novel Proteins of the Plasmodium Parasitophorous Vacuole Membrane. mSphere 3 (2018).

32. M. Zhang, S. Mishra, R. Sakthivel, B. M. Fontoura, V. Nussenzweig, UIS2: A Unique Phosphatase Required for the Development of Plasmodium Liver Stages. PLoS Pathog 12, e1005370 (2016).

33. T. Nessel et al., EXP1 is required for organisation of EXP2 in the intraerythrocytic malaria parasite vacuole. Cell Microbiol 22, e13168 (2020).

34. J. M. Przyborski et al., Trafficking of STEVOR to the Maurer’s clefts in Plasmodium falciparum-infected erythrocytes. EMBO J 24, 2306–2317 (2005).

35. A. S. Nasamu et al., Plasmepsins IX and X are essential and druggable mediators of malaria parasite egress and invasion. Science 358, 518–522 (2017).

36. P. Pino et al., A multistage antimalarial targets the plasmepsins IX and X essential for invasion and egress. Science 358, 522–528 (2017).

37. S. Bantuchai et al., Rhoptry neck protein 11 has crucial roles during malaria parasite sporozoite invasion of salivary glands and hepatocytes. Int J Parasitol 49, 725–735 (2019).

38. J. D. Bendtsen, H. Nielsen, G. von Heijne, S. Brunak, Improved prediction of signal peptides: SignalP 3.0. J Mol Biol 340, 783–795 (2004).

39. P. R. Gilson et al. (2023) Sequence elements within the PEXEL motif and its downstream region modulate PTEX dependent protein export in Plasmodium falciparum. (Authorea, Inc.).

40. M. Ressurreição, A. Fréville, C. Van Ooij (2023) Identification of a non-exported Plasmepsin V substrate that functions in the parasitophorous vacuole of malaria parasites. (Cold Spring Harbor Laboratory).

41. C. M. Ho et al., Malaria parasite translocon structure and mechanism of effector export. Nature 561, 70–75 (2018).

42. R. F. Waller, M. B. Reed, A. F. Cowman, G. I. McFadden, Protein trafficking to the plastid of Plasmodium falciparum is via the secretory pathway. EMBO J 19, 1794–1802 (2000).

43. A. Adisa et al., The signal sequence of exported protein-1 directs the green fluorescent protein to the parasitophorous vacuole of transfected malaria parasites. J Biol Chem 278, 6532–6542 (2003).

44. P. Favuzza et al., Dual Plasmepsin-Targeting Antimalarial Agents Disrupt Multiple Stages of the Malaria Parasite Life Cycle. Cell Host Microbe 27, 642–658 e612 (2020).

45. S. J. Tarr, A. R. Osborne, Experimental determination of the membrane topology of the Plasmodium protease Plasmepsin V. PLoS One 10, e0121786 (2015).

46. M. J. Coffey et al., An aspartyl protease defines a novel pathway for export of Toxoplasma proteins into the host cell. Elife 4 (2015).

47. A. Curt-Varesano, L. Braun, C. Ranquet, M. A. Hakimi, A. Bougdour, The aspartyl protease TgASP5 mediates the export of the Toxoplasma GRA16 and GRA24 effectors into host cells. Cell Microbiol 18, 151–167 (2016).

48. P. M. Hammoudi et al., Fundamental Roles of the Golgi-Associated Toxoplasma Aspartyl Protease, ASP5, at the Host-Parasite Interface. PLoS Pathog 11, e1005211 (2015).

49. C. H. Hsiao, N. Luisa Hiller, K. Haldar, L. J. Knoll, A HT/PEXEL motif in Toxoplasma dense granule proteins is a signal for protein cleavage but not export into the host cell. Traffic 14, 519–531 (2013).

50. M. J. Coffey et al., Aspartyl Protease 5 Matures Dense Granule Proteins That Reside at the Host-Parasite Interface in Toxoplasma gondii. Mbio 9 (2018).

51. W. J. Blakely, M. J. Holmes, G. Arrizabalaga, The Secreted Acid Phosphatase Domain-Containing GRA44 from Toxoplasma gondii Is Required for c-Myc Induction in Infected Cells. mSphere 5 (2020).

52. A. M. Cygan et al., Coimmunoprecipitation with MYR1 Identifies Three Additional Proteins within the Toxoplasma gondii Parasitophorous Vacuole Required for Translocation of Dense Granule Effectors into Host Cells. mSphere 5 (2020).

53. M. A. Fierro, T. Hussain, L. J. Campin, J. R. Beck (2022) Knock-sideways by inducible ER retrieval enables a novel approach for studying Plasmodium secreted proteins. (Cold Spring Harbor Laboratory).

54. M. Shea et al., A family of aspartic proteases and a novel, dynamic and cell-cycle-dependent protease localization in the secretory pathway of Toxoplasma gondii. Traffic 8, 1018–1034 (2007).

55. M. Franco et al., A Novel Secreted Protein, MYR1, Is Central to Toxoplasma’s Manipulation of Host Cells. mBio 7, e02231–02215 (2016).

56. N. D. Marino et al., Identification of a novel protein complex essential for effector translocation across the parasitophorous vacuole membrane of Toxoplasma gondii. PLoS Pathog 14, e1006828 (2018).

57. S. H. Adjalley, M. C. S. Lee, D. A. Fidock, “A Method for Rapid Genetic Integration into Plasmodium falciparum Utilizing Mycobacteriophage Bxb1 Integrase”. (Humana Press, 2010), 10.1007/978-1-60761-652-8_6, pp. 87-100.

58. M. Garten et al., EXP2 is a nutrient-permeable channel in the vacuolar membrane of Plasmodium and is essential for protein export via PTEX. Nat Microbiol 3, 1090–1098 (2018).

59. T. N. Petersen, S. Brunak, G. von Heijne, H. Nielsen, SignalP 4.0: discriminating signal peptides from transmembrane regions. Nat Methods 8, 785–786 (2011).

60. F. Sievers et al., Fast, scalable generation of high-quality protein multiple sequence alignments using Clustal Omega. Mol Syst Biol 7, 539 (2011).

61. A. M. Waterhouse, J. B. Procter, D. M. Martin, M. Clamp, G. J. Barton, Jalview Version 2--a multiple sequence alignment editor and analysis workbench. Bioinformatics 25, 1189-1191 (2009).

